# Unipept Desktop 2.0: construction of targeted reference protein databases for proteogenomics analyses

**DOI:** 10.1101/2023.02.09.527820

**Authors:** Pieter Verschaffelt, Alessandro Tanca, Marcello Abbondio, Tim Van Den Bossche, Tibo Vande Moortele, Peter Dawyndt, Lennart Martens, Bart Mesuere

**Affiliations:** Department of Applied Mathematics, Computer Science and Statistics, Ghent University, Ghent, Belgium; VIB - UGent Center for Medical Biotechnology, VIB, Ghent, Belgium; Department of Biomedical Sciences, University of Sassari, Sassari, Italy; Department of Biomolecular Medicine, Faculty of Medicine and Health Sciences, Ghent University, Ghent, Belgium

## Abstract

Unipept Desktop 2.0 is the most recent iteration of the Unipept Desktop tool that adds support for the analysis of proteogenomics datasets. Unipept Desktop now supports the automatic construction of targeted protein reference databases that only contain proteins associated with a predetermined list of taxa. This improves both the taxonomic and functional resolution of a metaproteomic analysis and yields several technical advantages. By limiting the proteins present in a reference database, it is now also possible to perform (meta)proteogenomics analyses. Since the protein reference database now lives on the user’s local machine, they have complete control over the database used during an analysis. Data does no longer need to be transmitted over the internet, decreasing the time required for an analysis and better safeguarding privacy sensitive data. As a proof of concept, we present a case study in which a human gut metaproteome dataset is analyzed with Unipept Desktop 2.0 using different targeted databases based on matched 16S rRNA gene sequencing data.

## Introduction

The metaproteomics research discipline has undergone a big transition since the term was first introduced in 2004^1^. We have witnessed the evolution of metaproteomics from very small-scale experiments, in which three distinct proteins^2^ could be identified in an ecosystem, to a mature technology that is able to analyze more than 100 000 protein fragments (or peptides) from various environments.

Unipept^3^ was one of the first major tools in this promising new research field that could be used to analyze tryptic peptide-based metaproteomics samples. Originally starting as a web application, Unipept was quickly accompanied by an application programming interface^4^ (API) and command line interface^5^ (CLI) that respectively allow for embedding Unipept’s analyses in other tools and analyzing larger samples directly from the command line. API usage metrics currently indicate that more than 500 000 requests are handled by Unipept on a monthly basis, acknowledging the importance of the tool in this field.

The advent of recent technological improvements in mass spectrometry and more powerful proteomics approaches have allowed metaproteomics to transition from small studies to large scale experiments^6,7^. Due to Unipept’s inherent web-based nature, it was limited in the size of the samples that could be analyzed because of browser-imposed restrictions on available compute resources. This led to the development of the Unipept Desktop application^8^ in 2020.

Unipept Desktop^8^ paved the way for the analysis of large metaproteomics samples (containing 500k peptides or more) and completely overhauled the way these metaproteomics samples can be organized with the introduction of projects and studies. Projects can easily be shared with other researchers, who no longer need to reanalyze samples and wait for the results to become available, or can be archived for later use. The Unipept Desktop application also introduces a new inter- and intra-sample comparison pipeline which allows users to gain insight into the taxonomic and functional shift within and between multiple samples.

Over the last few years, interest in a new research area, proteogenomics, has been growing. Proteogenomics can be thought of as a logical next step in researching complex microbial ecosystems and combines information from both metagenomics and metaproteomics experiments in order to overcome a few key problems that arise when working with metaproteomics data in isolation^9^.

The first major issue that we need to consider is the ever-growing size of the protein reference databases that are being used to match peptides with proteins. UniProtKB^10,11^, a freely accessible database containing protein sequences, has seen a rapid increase in size over the last decade and has grown from approximately 19 million proteins in 2012 to 227 million proteins in 2022. Compared to the early days of Unipept, we are able to identify increasingly diverse species as a direct result of the increased size of the reference database, but this also comes with a few drawbacks. Each peptide that is presented to Unipept will be matched with all proteins in which this peptide occurs. All of these proteins are associated with a specific organism and such a peptide-based analysis thus results in a set of organisms from which this peptide could potentially originate. In order to increase insight of researchers into the taxonomic composition of a sample, Unipept summarizes all of this information and calculates the lowest common ancestor (LCA) of this set of organisms (i.e., NCBI taxa) for each peptide. If all of the matched organisms are evolutionarily close to each other, this works very well and the LCA of our matches will be of value. If, however, one or more of the matched organisms is very different from the others, the LCA will typically end up at the root or another very general taxon within the NCBI taxonomy (Figure 1a).

**Figure 1:**
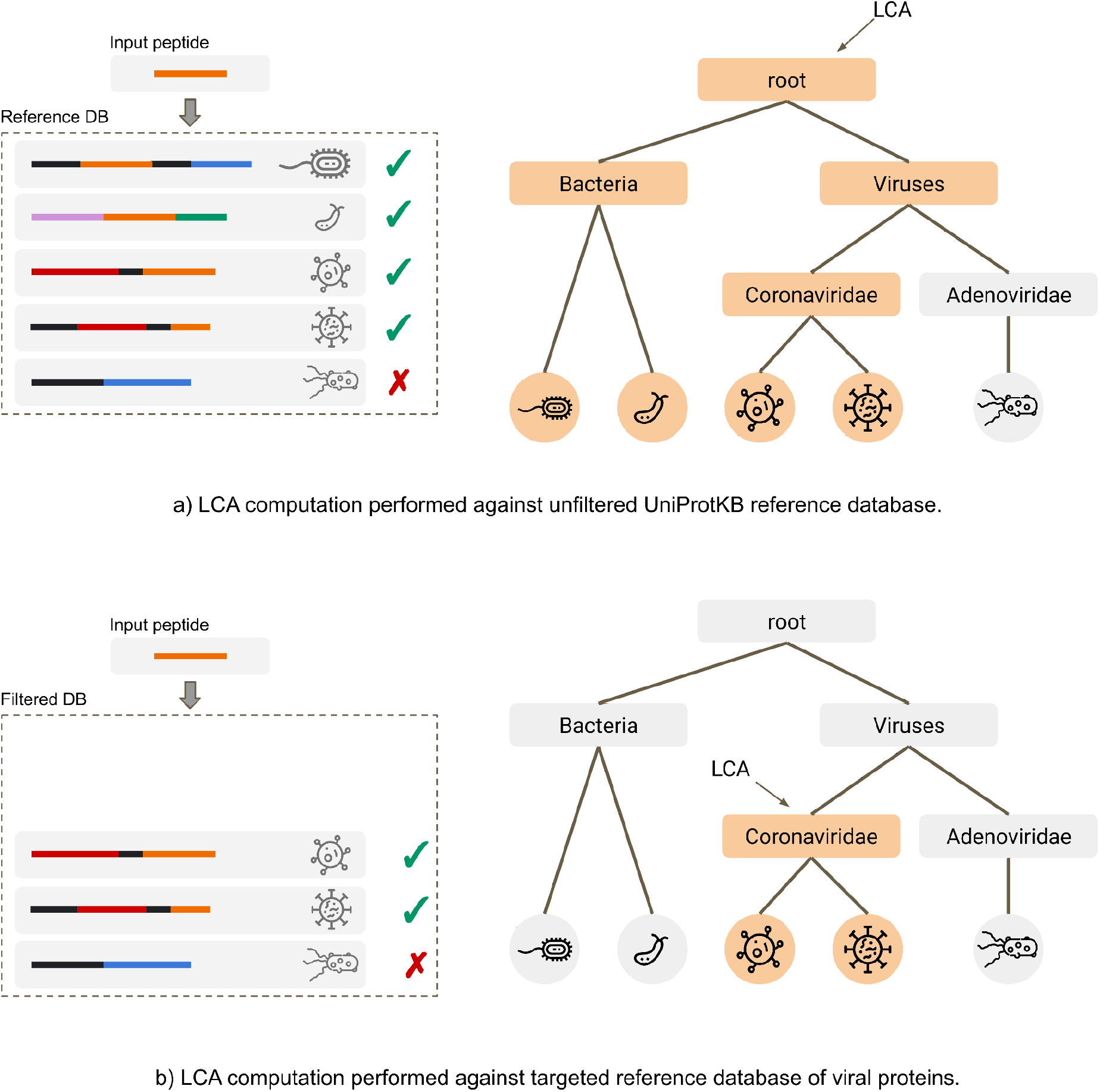
Two examples of how the lowest common ancestor (LCA) for a set of identified taxa is computed in Unipept Desktop. In both cases, a single input peptide is matched against proteins in a reference database. The taxa associated with all matched proteins are then summarized as the LCA, which is the most specific node in the taxonomy tree that is a parent of all matched taxa. An unfiltered protein reference database is used in the matching process. Since more proteins from a more diverse range of species are matched, the LCA ends up at the root of the taxonomy tree, providing little to no information. **b)** The reference database is restricted to viral proteins only and a much more specific LCA will be found (mapping onto the *Coronaviridae* family). These examples illustrate the importance of targeted protein reference databases when analyzing metaproteomics samples.

Proteogenomics tries to overcome this problem by combining information from prior metagenomics experiments from the same environment with metaproteomics experiments. The metagenomics experiment is used to explore the taxonomic composition of an ecosystem, which subsequently guides the researcher to query only a subset of the reference database. In the case of shotgun metagenomics, DNA sequences identified by a metagenomics experiment can be used to build a customized protein reference database^12^.

When using the Unipept Web application, analyses are always performed against the entire UniProtKB resource. We previously added the possibility to set up a local instance of Unipept and search against the database provided by this endpoint as part of the Unipept Desktop application, but setting up such a custom endpoint is often experienced as a big technical hurdle by our target audience. Most researchers are thus still dependent on the reference database provided by Unipept, including the database update scheme that Unipept dictates. This creates a set of problems and drawbacks that need to be overcome in order to properly support proteogenomics data analyses. Since users do not have control over the database that is being used, they cannot provide potential metagenomics information (such as taxa identified in the ecosystem under study) and restrict the search space of the reference database.

To solve this problem, we introduce version 2.0 of the Unipept Desktop application, which marks the beginning of a new era for the analysis of proteogenomics datasets. Unipept Desktop now provides support for the automatic construction of targeted protein reference databases on the user’s local machine. Such targeted databases are based on a filtered version of UniProtKB and only contain UniProtKB records that are associated with the taxa provided by the user (Figure 1b).

In this article, we discuss how these targeted protein reference databases are constructed by the Unipept Desktop application and how they can be queried efficiently on a user’s machine. We also present a case study, based on a human gut metaproteome dataset obtained from 28 celiac patients, in which we investigate to what extent the accuracy of the analysis results improves by only taking into account a subset of the UniProtKB reference database.

## Unipept Desktop 2.0

In order to perform a proteogenomics experiment, researchers typically first perform a metagenomics experiment on the environment of interest, which then provides them with a set of organisms that are likely to be present in that environment. 16S rRNA gene sequencing and shotgun metagenomics are two widely used techniques in this research area that provide the required information, but other types of analysis can also be used. Sometimes the taxonomic composition of an environment can also be inferred from previous studies, if available.

Once the approximate taxonomic composition of a specific sample is known, a targeted protein reference database can be created that contains only those proteins that can be produced by the organisms of interest. This helps to drastically reduce the search space for a subsequent metaproteomics analysis (typically performed on the same environment) that is performed using the newly constructed targeted protein reference database.

Version 2.0 of the Unipept Desktop application is fully focused on the analysis of these proteogenomics datasets and therefore introduces an exciting new feature that allows users to construct targeted protein reference databases in two different ways.

To construct a targeted protein reference database with Unipept Desktop 2.0, researchers need some way of selectingUniProtKB proteins that can be present in the final result. The first selection method allows to specify which proteins are retained by providing a list of valid NCBI taxon identifiers. Unipept will then only include those proteins in the final database that are associated with a taxon that is present in this list, or that are associated with an (in)direct child of one of the provided taxon identifiers. Note that there’s also the option to limit the database construction process to SwissProt (instead of SwissProt + TrEMBL) such that only manually-curated proteins are included. The second selection method requires the user to provide a set of UniProtKB reference proteome identifiers. Only the proteins that are found in these reference proteomes will then be selected for the construction of the reference database.

### Construction of targeted protein reference databases

Protein reference databases that are required for the analysis of metaproteomic samples are typically very large. Depending on the number of proteins included, the size of these databases ranges from a few gigabytes to more than a terabyte. It is these huge size requirements that make it technically very hard to build targeted protein reference databases for multiple concurrent users on Unipept’s servers.

Moving protein reference databases from Unipept’s servers to a researcher’s local machine opens up a world of new possibilities.

Firstly, researchers now have complete control over the database to be used for an analysis. They are no longer dependent on the update schedule dictated by Unipept, but can update their local reference database whenever they see fit. Previously, there was no way to roll back the database to the previous iteration of the UniProtKB resource after an update had been performed on Unipept’s servers. This is important because researchers could no longer correctly compare metaproteomics analysis results for samples that were analyzed using a different database version.

Secondly, privacy sensitive data, which may be part of some metaproteomics experiments, is no longer sent over the internet to the Unipept servers for analysis. Some research institutes or applications do not allow sensitive data to be sent to remote services for analysis, but rather require it to be kept in-house to protect patient confidentiality and privacy.

Thirdly, in most cases, the runtime of analyses performed using a local database is improved because the data no longer needs to be transmitted over the Internet and the targeted protein reference databases are typically much smaller in size. The smaller the reference database, the faster the analyses can be performed.

Finally, the amount of false positive peptide matches that can occur is drastically reduced when compared to metaproteomics analyses against the complete UniProt database.

### Implementation

#### Dependency management and portability

Unipept uses a custom format for storing protein reference databases, which includes a large amount of pre-computed data to speed up subsequent metaproteomics analyses. This database format is loaded into a relational database management system (RDBMS) such as MySQL. An RDBMS is a very specialized piece of software designed to query huge amounts of structured data as quickly as possible.

The Unipept Desktop application does not query this database directly, but instead relies on the Unipept API to do the hard work. The Unipept API provides a standardized set of HTTP REST endpoints that respond to queries from the Unipept Desktop (or Unipept Web) application with the desired information. This Unipept API in turn is a Ruby-on-Rails project that needs to be executed by a piece of software called a web server.

Both the installation and configuration of an RDBMS and a web server (to run the Unipept API) require a considerable amount of time, effort and technical skills, which is undesirable for users of the Unipept Desktop application. They need an application that is easy to install and that does not require a lot of user intervention to start and maintain.

To address these issues, we decided to encapsulate all required software dependencies in a Docker^13^ image. Docker is a free tool that allows to run predefined virtual containers (as defined by an image) on a variety of different operating systems. A Docker image contains a set of instructions to be executed by a virtual computing environment (controlled by Docker) and guarantees that these instructions will work deterministically on any supported system. By relying on Docker for the RDBMS and the web server, we have reduced the number of dependencies that are required by the Unipept Desktop to just one: Docker (which can be downloaded for free from its official website and supports all major operating systems).

Communication between the Unipept Desktop app and the software dependencies that are managed by Docker is completely transparent to the user. We use a NodeJS package called Dockerode, which handles all communication between both parties.

#### Filtering UniProtKB by NCBI taxon identifiers

We described earlier how a targeted protein reference database can be constructed by selecting a list of NCBI taxon identifiers. In this case, Unipept selects only those proteins from the UniProtKB resource that are associated with one of these taxa (or children of these taxa) and includes them in the targeted reference database. In this section, we describe how this is implemented so that efficient filtering can be performed on all 227 million UniProtKB proteins.

The first step in the database construction process is downloading and processing all proteins from the UniProtKB database sources. For each source (SwissProt and TrEMBL), the application constructs a database index structure that can be easily queried and reused in the future. This index structure consists of several “chunks”, each containing a set of different proteins. These chunks are compressed line-based ‘tsv’-files that contain all the necessary information (protein identifier, linked NCBI taxon identifiers, functional annotations, etc.). The protein information is sorted numerically by NCBI taxon identifier, and each chunk contains only those proteins that correspond to a known subset of the NCBI taxon identifier space. Figure 3 shows a schematic representation of how this index structure is constructed and the impact it has on the final targeted database construction process.

**Figure 2:**
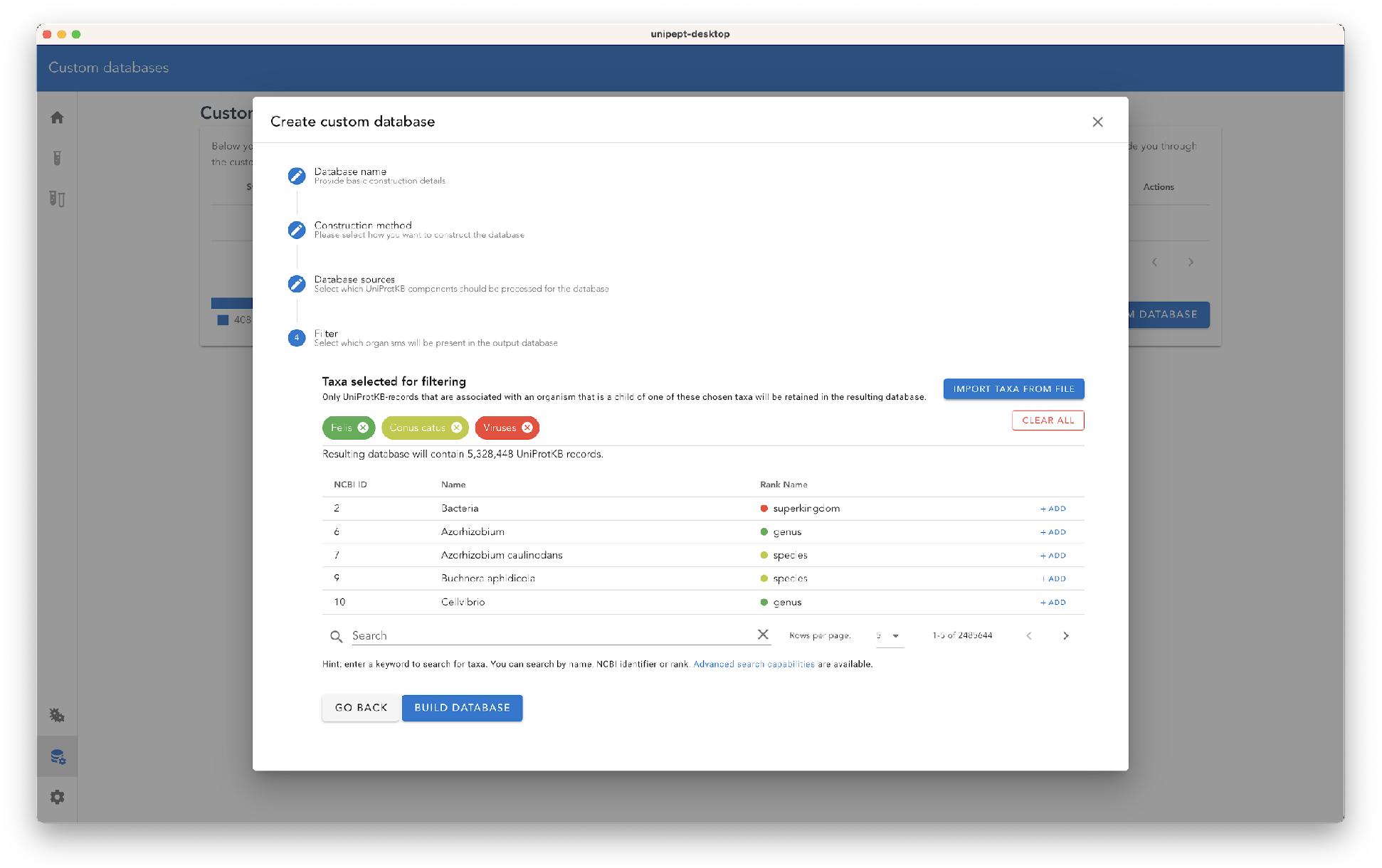
Database creation wizard in Unipept Desktop 2.0. This wizard guides the user through the process of building a targeted protein reference database. Researchers can select proteins in two different ways: by providing a set of taxon identifiers, or by providing a list of UniProtKB reference proteome IDs.

**Figure 3:**
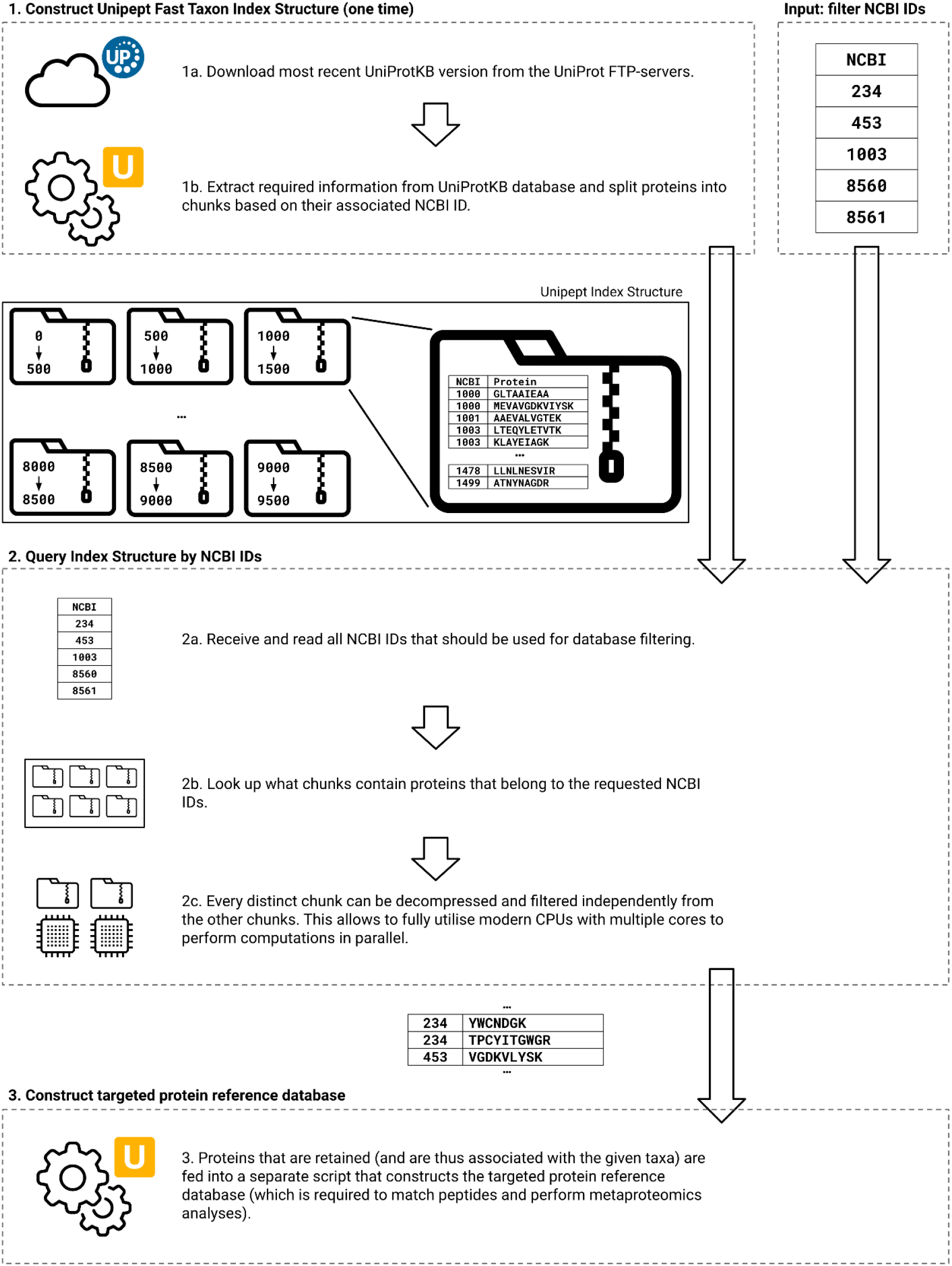
To efficiently filter the UniProtKB database, Unipept builds a custom index structure that can be easily filtered by taxon ID. This index structure only needs to be built the first time a targeted protein reference database is constructed. The index will be reused for all subsequent database builds. The NCBI IDs, which can optionally be provided by a user, are used to select which index chunks need to be queried. Each of these chunks is a compressed file containing protein information, and each of these chunks can be processed in parallel by multiple CPU-cores (speeding up the construction process).

Downloading and constructing the reusable database index structure is a process that only needs to be performed once for each version of the UniProtKB resource. Subsequent targeted protein reference databases will reuse an existing index or will automatically rebuild it if the UniProtKB resource has been updated since the previous time the index was constructed.

To efficiently query the index structure, we first determine which chunks need to be queried. This is simply a matter of looking up which ranges the various NCBI IDs provided fall into and only processing the chunks that correspond to those taxon ranges. All of these chunks are completely disjoint from each other and can be processed and filtered in parallel, maximizing the use of modern multi-core CPUs.

## Case Study

To assess the strength of proteogenomics analyses in Unipept Desktop, we used Unipept Desktop 2.0 to perform the taxonomic annotation of a metaproteomic dataset obtained from 28 human fecal samples collected from celiac disease patients following a gluten-free diet and previously subjected to a 16S rRNA gene sequencing study^14^. Here, we re-analyzed and re-annotated the 16S rRNA gene sequencing data using a robust and up-to-date bioinformatics pipeline based on the amplicon sequence variant (ASV) approach and a newer database^15^ (as previously described^16^), to obtain accurate information about the set of bacterial taxa present in the environment under study. In parallel, the residues from the 28 fecal samples underwent protein extraction, filter-aided sample preparation and LC-MS/MS analysis, according to established procedures^17^. Mass spectra were analyzed by Proteome Discoverer (version 2.4, Thermo Fisher Scientific)^18^, using a publicly available collection of human gut metagenomes^19^ as the sequence database, Sequest-HT as the search engine and Percolator for peptide validation (search parameters are detailed elsewhere). A total of 64 845 microbial peptides (of which 62 363 peptides remained after duplicate filtering) were identified (with 1% as FDR threshold) and used as input for Unipept Desktop annotation. We used the online NCBI Taxonomy Browser^20^ to convert between taxonomy IDs and taxa names where necessary.

The purpose of this case study was to demonstrate that the Unipept Desktop app is capable of analyzing a metaproteomic sample by matching the input peptides to only a specific subset of the proteins in the UniProtKB^21^ database. Secondly, we investigated the extent to which the taxonomic profile of a metaproteomic dataset differs when annotated against different protein reference databases.

As the 16S rRNA gene is only present in bacterial species, we restricted our analysis to bacteria. All metaproteomic analyses in this study are performed using Unipept Desktop v2.0.0-alpha.7 with the search settings “filter duplicates” and “equate I/L” enabled (the “advanced missed cleavage handling” setting was disabled).

The experiment started by processing all 227 million proteins in UniProtKB 2022.3 (both SwissProt and TrEMBL) and constructing an initial, unfiltered, protein reference database. During the analysis of our metaproteomic sample using Unipept Desktop and this general-purpose database, we were able to match 56 070 peptides (out of a total of 62 363 peptides, or 89.9%). Of these matches, 30 248 peptides (53.9%) were annotated with a taxon at the rank of “family” (or lower) and 13 659 peptides (24.4%) were annotated with a taxon at the rank of “species” (or lower).

We repeated this analysis using three other targeted protein reference databases, constructed by including only all taxa identified by the 16S rRNA gene sequencing analysis at the family, genus or species ranks, respectively. The higher a rank in the NCBI taxonomy, the less specific the constructed reference databases will be and the more proteins they will still contain. For each NCBI rank of interest, we counted the number of peptides annotated with a taxon of that rank (or lower) and compared these counts across the different protein reference databases. As can be seen in Figure 4, the total number of peptides matched decreases as the size of the protein reference databases decreases, but the number of taxon matches at a lower rank of the NCBI taxonomy increases for all three targeted protein reference databases.

**Figure 4:**
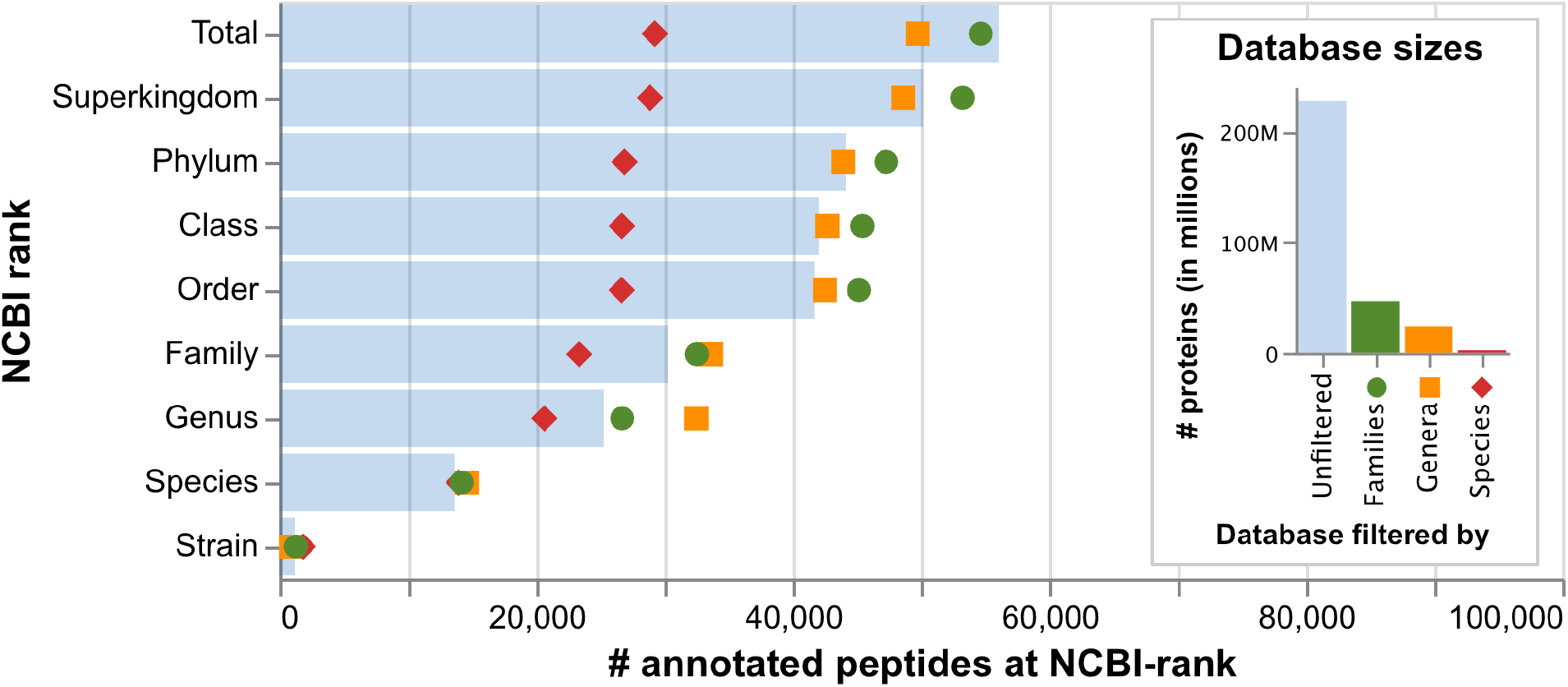
Four analyses have been performed on the same metaproteomic sample, but with a different underlying protein reference database. In the main visualization (left), we compare the number of peptides with a taxon match at a specific rank (or lower) in the NCBI taxonomy. In most cases the analyses performed by using a reference database constructed from families or genera performs as well as or better than the full UniProtKB database. However, these databases contain only a fraction of the proteins (right) and therefore require much less computational resources. At the species level, all protein reference databases behave almost identically. At higher ranks, we can see that the reference database constructed from a set of species is probably too restricted.

Looking at the higher ranks, we see that the larger reference databases have an advantage and provide a valuable annotation for more peptides (53 243 (95.0%), 48 637 (86.7%) and 28 877 (51.5%) taxon matches at the “Superkingdom” rank or lower for the “families”, “genera” and “species” based databases, respectively). Note that, at this level, the targeted database constructed from a list of “families” already outperforms the unfiltered database (Figure 4).

It is important to note that there is a trade-off to be made between matches on a particular NCBI rank of interest and the input filter used to construct targeted protein reference databases. The more restrictive the input filter (assuming it is a good representation of the taxa in the environment under study), the fewer peptides will generally be matched, but the more likely it is that more detailed taxon matches will be found. This is easily explained. The more proteins there are in a reference database, the greater the chance that a purely random peptide match will occur by chance. These random matches are typically with proteins from organisms unrelated to the environment of interest, so the lowest common ancestor calculation ends up at a very high level in the NCBI taxonomy (Figure 1).

Going down a few ranks in the NCBI taxonomy and counting the number of matches at the species level, we find 13 659 taxa (24.4%) when using the unfiltered database and 14 192 (25.3%), 14 593 (26.0%) and 13 960 matches (24.9%) for the “families”, “genera” and “species” based databases, respectively. At this level, all three of the targeted reference databases outperform the unfiltered database while containing significantly fewer proteins. These significantly smaller protein reference databases require less storage space and fewer computing resources to work with, allowing the analyses to be performed on a simple laptop (rather than relying on the remote Unipept web servers). In most cases, the analysis is completed faster (especially when the missed cleavage handling option is enabled) and the end-user has full control over exactly which database is being used.

If we compare the targeted analysis with the analysis based on the unfiltered database, we see that 1 408 peptides that were previously annotated are now unmatched. As this is not an insignificant amount, it is important to investigate what the main differences between the two analyses at a taxonomic level are.

If we look at the taxa that were matched using the full database, and not when using the targeted database, we see that most of them belong to the *Firmicutes* phylum (813 matches). Looking a little further, we see that the species *Evtepia gabavorous* (80 matches), *Odoribacter splanchnicus* (44 matches), and *Turicibacter sanguinis* (34 matches) are the most represented within the *Firmicutes*.

According to the NCBI taxonomy, *Evtepia gabavorous* belongs to the *Eubacteriales incertae sedis* “family”, which is actually an unclassified taxon and is therefore not included in the 16S SILVA database. Secondly, *Odoribacter splanchnicus* belongs to the *Marinifilaceae* family according to SILVA, whereas it is assigned to the *Odoribacteraceae* family by the NCBI taxonomy (which is why this family is not present in the taxon list that was used to construct a protein reference database and is therefore not matched during the Unipept analysis). Finally, a similar explanation applies to*Turicibacter*, which belongs to *Erysipelotrichaceae* according to SILVA and to the *Turicibacteraceae* family according to NCBI. Therefore, this family is also missing from the filtered taxon list and its proteins are not included in the database construction.

Most of the other remaining *Firmicutes* assignments are mainly attributed to *Clostridiales* bacterium and *Firmicutes* bacterium (413 matches in total), two taxa for which the rank is *unspecified* in NCBI and which are therefore not taken into account when constructing the targeted database. A further 122 peptide matches using the full database were taxonomically annotated at the root level, which does not provide any valuable information other than the fact that the peptide was indeed matched.

After this detailed analysis, we can conclude that most of the mismatches when using a targeted reference database are due to taxonomic inconsistencies between the SILVA database (used for the 16S rRNA gene taxonomic annotation) and the NCBI taxonomy that is being used by Unipept, or even to incomplete or provisional annotations within the NCBI taxonomy itself. This problem could be overcome by also using NCBI for the taxonomic classification of 16S data (instead of SILVA), but this choice is entirely up to the end-user, and is outside the scope of the analysis performed by Unipept Desktop.

All of these experiments have been conducted on a normal modern computer with a 6-core CPU (AMD Ryzen 3600X),16GiB of RAM, a SATA-6 SSD and a 100Mbps internet connection. On this machine, it took approximately 21 hours and 25 minutes to construct a custom database containing 46 million proteins or approximately 20 minutes for a targeted database containing 1 million proteins. All proteins from the UniProtKB resource are preprocessed the first time a targeted database is constructed and took approximately 5 hours and 30 minutes on this machine (this preprocessing step will automatically be performed when the UniProtKB resource updates). The final size of the database cache is 52GiB, and the 46 million and 1 million protein databases are respectively 255GiB and 5.76GiB in size.

## Concluding remarks

The newest iteration of the Unipept Desktop app builds upon the strength of the existing Unipept infrastructure to enable support for the analysis of proteogenomics samples. Leveraging taxonomic information of the environment under study (e.g. generated from a metagenomics experiment), it is possible to construct targeted protein reference databases that include only a (relevant) subset of proteins from the UniProtKB resource. These significantly smaller reference databases drastically improves the time and computational resources required to subsequently analyze metaproteomics samples, which ultimately makes it possible to perform these analyses on a local machine.

Since Unipept Desktop 2.0 makes it possible to perform metaproteomics analysis on a local machine, a range of new possibilities opens up. Privacy-sensitive data no longer needs to be transmitted over the internet and users now control which reference database is used. We have shown that using targeted protein reference databases can even lead to a metaproteomics analysis with a higher taxonomic resolution (assuming that the selected taxa suits the environment under study).

No scientific software package is ever completed and we can still think about future improvements that could be beneficial for the Unipept Desktop application. First of all, at this point it is not yet possible to construct targeted protein reference databases from UniProtKB versions other than the current one. This is a consequence of the fact that previous versions of the UniProtKB database are provided in a different file format for which a new parser needs to be implemented.

Secondly, since all targeted protein reference databases are always constructed by filtering the UniProtKB resource, only proteins that are included in UniProtKB can be matched using Unipept. This can be problematic for some research disciplines (such as protein research of ocean water) that are investigating proteins of organisms that are not well represented in UniProtKB. This problem could be overcome by allowing Unipept to construct protein reference databases from external sources (e.g., represented by FASTA or PEFF files). These additions are considered for future versions of the Unipept Desktop app.

## Acknowledgements

This work has benefited from collaborations facilitated by the Metaproteomics Initiative (https://metaproteomics.org/) whose goals are to promote, improve, and standardize metaproteomics^22^. This work has furthermore been supported by the Research Foundation - Flanders (FWO) [1164420N to P.V., 12I5220N to B.M.] and by the University of Sassari [Fondo di Ateneo per la Ricerca 2020 to A.T.]. We thank the Flemish Supercomputer Center (VSC) funded by the Research Foundation - Flanders (FWO) and the Flemish Government for providing the infrastructure to build the Unipept database and to run the experiments from this manuscript. Part of this work was also supported by the Research Foundation - Flanders (FWO) for ELIXIR Belgium [I002819N].

